# The chaperonin GroESL facilitates *Caulobacter crescentus* cell division by supporting the function of the actin homologue FtsA

**DOI:** 10.1101/2020.12.17.423375

**Authors:** Kristen Schroeder, Kristina Heinrich, Ines Neuwirth, Kristina Jonas

## Abstract

The highly conserved chaperonin GroESL performs a crucial role in protein folding, however the essential cellular pathways that rely on this chaperone are underexplored. Loss of GroESL leads to severe septation defects in diverse bacteria, suggesting the folding function of GroESL may be integrated with the bacterial cell cycle at the point of cell division. Here, we describe new connections between GroESL and the bacterial cell cycle, using the model organism *Caulobacter crescentus*. Using a proteomics approach, we identify candidate GroESL client proteins that become insoluble or are degraded specifically when GroESL folding is insufficient, revealing several essential proteins that participate in cell division and peptidoglycan biosynthesis. We demonstrate that other cell cycle events such as DNA replication and chromosome segregation are able to continue when GroESL folding is insufficient, and find that deficiency of the bacterial actin homologue FtsA function mediates the GroESL-dependent block in cell division. Our data suggest that a GroESL-FtsA interaction is required to maintain normal dynamics of the FtsZ scaffold and divisome functionality in *C. crescentus*. In addition to supporting FtsA function, we show that GroESL is required to maintain the flow of peptidoglycan precursors into the growing cell wall. Linking a chaperone to cell division may be a conserved way to coordinate environmental and internal cues that signal when it is safe to divide.

**Importance:** All organisms depend on mechanisms that protect proteins from misfolding and aggregation. GroESL is a highly conserved molecular chaperone that functions to prevent protein aggregation in organisms ranging from bacteria to humans. Despite detailed biochemical understanding of GroESL function, the *in vivo* pathways that strictly depend on this chaperone remain poorly defined in most species. This study provides new insights into how GroESL is linked to the bacterial cell division machinery, a crucial target of current and future antimicrobial agents. We identify a functional interaction between GroESL and FtsA, a conserved bacterial actin homologue, suggesting that as in eukaryotes, some bacteria exhibit a connection between cytoskeletal actin proteins and chaperonins. Our work further defines how GroESL is integrated with cell wall synthesis, and illustrates how highly conserved folding machines ensure the functioning of fundamental cellular processes during stress.

## Introduction

All life must monitor and adjust the vital processes of growth and division in response to external and internal environmental cues. Prokaryotic model organisms offer an accessible system to study how essential biological processes are regulated in response to these cellular and environmental signals. Molecular chaperones are ubiquitous proteins with high sequence conservation, and studying protein folding dynamics in bacterial systems has been crucial in understanding the fundamental processes that assist all organisms in building and maintaining functional proteins.

During biosynthesis, some proteins must overcome energy barriers in order to achieve their native fold (1). For these proteins, interaction with ATP-powered chaperones assists them in attaining a functional conformation on a biologically relevant time scale *in vivo* (1). This chaperone interaction is not limited to biosynthesis, as changes in intracellular conditions, for example temperature or oxidative stress, or the presence of toxic compounds, can destabilize folding of a wide array of proteins (2–4), which may then require refolding. To adjust chaperone folding capacity to these different folding demands, the expression of chaperone genes can be increased above basal levels through stress-responsive transcriptional control, for example through induction by the heat shock sigma factor (5). In this way, chaperone folding capacity is available for synthetic processes during optimal conditions, and increased to rescue misfolding proteins during diverse stresses.

The majority of ATP-powered protein folding in prokaryotes is carried out by the highly conserved DnaK/J/GrpE-ClpB bichaperone system and the GroES/EL chaperonin machine (1), which are assisted in interacting with their client proteins by a network of less-conserved holdases, including small heat shock proteins and chaperedoxins (6). GroEL (Hsp60, Cpn60) is a heat shock protein that oligomerizes into a tetradecameric double ring structure with two central cavities that can capture unfolded proteins via their solvent-exposed hydrophobic residues (7). The GroES (Hsp10, Cpn10) co-chaperonin then binds as a lid over the GroEL-client complex, encapsulating client proteins and thus providing a segregated environment to assist with folding (7). While the number and arrangement of the *groES groEL* genes varies across bacteria, they are most often found in a single copy together in an operon that allows both for housekeeping expression, for example from a *σ*^70^-dependent promoter, as well as stress-responsive expression, for example from a *σ*^32^-dependent promoter or HrcA repressor sequences (5, 8, 9).

Despite detailed description of GroESL folding mechanics and a good understanding of the regulation of *groESL* transcription, comparably few studies exist examining the role of chaperonins in physiological processes. With the exception of some Mollicutes (10), GroESL is essential in all bacterial species investigated to date (9, 11–14). Several of these organisms exhibit a relationship between chaperonin availability and the cell cycle, as cell division is blocked when GroESL levels are reduced (13–15). During bacterial cell division the presence, location, and quantity of many different classes of proteins, as well as the remodelling of the cell envelope must be tightly controlled in order to successfully make two daughter cells. The regulation of these proteins can occur at the level of synthesis, degradation, conformation, localization and activity (16, 17), however the contribution of protein folding state and chaperone interactions to cell division is not yet well understood.

The most well-studied bacterial chaperonin is that of *Escherichia coli*, where ∼250 proteins have been identified as interacting partners of GroESL (18, 19). Of these client proteins, 57 have been shown to obligately depend on GroESL for folding into the native state (classified as obligate, or Type IV GroESL substrates), including 6 essential proteins (18–20). One of the identified essential obligate GroESL substrates is the cell division protein FtsE (19), however while a functional deficit of this protein may contribute to the cell division defect reported during GroESL depletion in *E. coli* (21), the requirement for FtsE function can be bypassed by altering growth conditions, and it therefore remains unclear if FtsE is conditionally linked to GroESL in this organism. Chaperonin studies in other bacteria have identified a few proteins from the GroESL client protein pool (22–24), and it remains poorly understood how GroESL function impacts the cell cycle and other physiological processes in these organisms.

The model organism *Caulobacter crescentus* is a Gram-negative oligotrophic alpha-proteobacterium with a dimorphic lifestyle that produces two morphologically distinct daughter cells (25). This asymmetric life cycle has made *Caulobacter* a powerful model for investigating events of the cell cycle. In particular, cell division has been well described in this organism, which has led to important advances being made in understanding the conserved mechanisms mediating cell division in bacteria (26, 27). Similarly to other bacteria, *C. crescentus* becomes filamentous when GroESL is depleted (15), indicating an involvement of GroESL in the cell cycle. However, the precise role of GroESL in *Caulobacter* cell cycle progression and cell division has not been studied so far.

In this study, we establish the connection between GroESL folding and cell division in *C. crescentus*. We have identified a subset of the proteome that changes solubility depending on the presence of the chaperonin, and show that several proteins involved in cell division and peptidoglycan (PG) biosynthesis become insoluble when GroESL folding capacity is reduced.

Furthermore, we find that the bacterial actin homologue FtsA is responsible for mediating the filamentation phenotype observed when GroESL-mediated folding is insufficient, suggesting a relationship between chaperonins and actin-like proteins in prokaryotes. Integrating chaperonin folding into cell division in this way may represent a way of coordinating environmental and internal cues that signal when it is safe to divide.

## Results

### GroESL folding insufficiency results in filamentation

As GroESL is essential in *Caulobacter* and cannot be deleted (12), we made use of the previously described *C. crescentus* strain SG300, where the regulatory region upstream of *groESL* is replaced by a xylose-inducible promoter (Figure 1A) (12). When xylose is removed from the growth media, GroESL is diluted from the growing culture over several hours (Figure 1B). As GroESL levels decline, the demand for chaperonin-mediated folding exceeds what can be provided, resulting in the development of phenotypes associated with client protein misfolding (Figure 1C). In *C. crescentus*, cells grow into long filaments featuring wide segments interspersed with irregular shallow constrictions (Figure 1C) (15). When GroESL levels fall to 30% of the wild type (4h depletion, Figure 1D), cell lengths diverge from normal population lengths, and the shallow constrictions are properly localized at midcell (Figure 1C).

**Figure 1:**
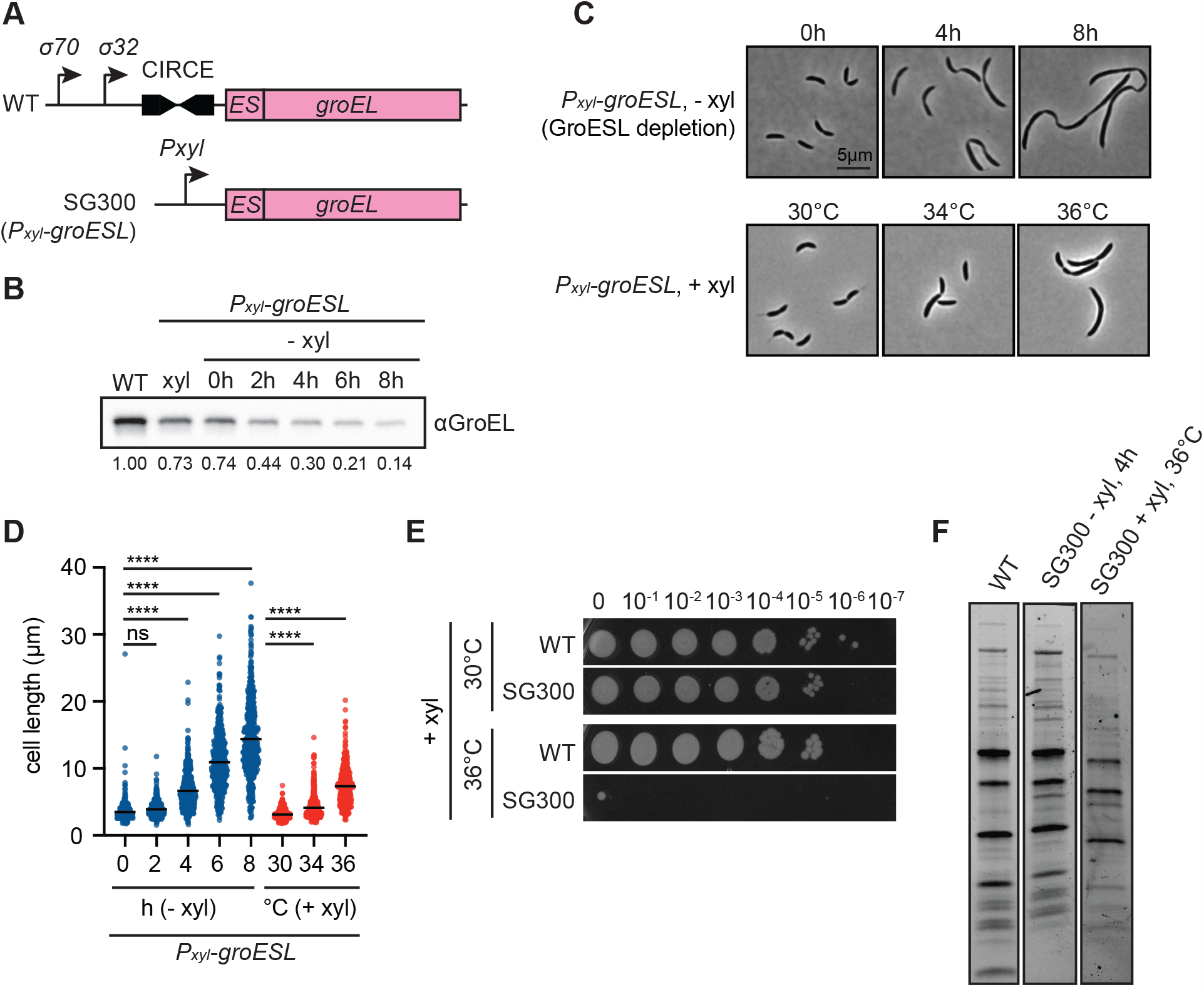
GroESL folding insufficiency results in cell filamentation. (A) Diagram of GroESL locus in wild type (WT) and GroESL depletion (SG300, *P*_*xyl*_*-groESL*) strains. Wild type GroESL is regulated by the CIRCE element as well as a sigma 32-dependent promoter, which has been replaced by the xylose-inducible promoter in strain SG300 (12). (B) Western blot of protein levels of GroESL during depletion (-xyl). Quantifications of band intensities are an average of three biological replicates. (C) Phase contrast microscopy showing the morphology of the GroESL depletion strain (*P*_*xyl*_*-groESL*) when grown in depleting conditions (-xyl), in non-depleting conditions (+xyl), and at increased temperature. Cultures were depleted for the indicated time, or incubated for four hours at the indicated temperatures prior to fixation and imaging. (D) Quantification of population cell lengths of the GroESL depletion strain (*P*_*xyl*_*-groESL*) during depletion (-xyl) or when grown under non-depleting conditions (+xyl) at the indicated temperatures for four hours (n < 417 each population; ****, p < 0.0001). (E) Spot assay of wild type (WT) and the GroESL depletion strain (SG300), when grown in non-depleting conditions, incubated at the indicated growth temperatures. Xylose was included in all agar plates; plates were incubated for 2-3 days prior to imaging. (F) Coomassie staining of SDS-PAGE gel of insoluble fractions isolated from wild type and the GroESL depletion strain (SG300), grown either for four hours in depleting conditions (-xyl), or incubated at 36C for four hours in non-depleting conditions (+ xyl).

As protein folding is destabilized by temperature stress, we also investigated the heat sensitivity of the GroESL depletion strain, which at maximal induction produces less GroESL (73%) than the wild type (Figure 1B), and is unable to upregulate *groESL* transcription during stress conditions. At the optimal growth temperature of 30°C, no difference in viability was observed between wild type *C. crescentus* and the GroESL depletion strain grown in the presence of xylose (Figure 1E). However, a mild temperature increase to 36°C caused cultures of the GroESL depletion strain to filament and become inviable (Figure 1C, D), emphasizing the importance of upregulating GroESL to provide chaperonin-mediated folding at elevated temperatures. Wild type *C. crescentus* also exhibits filamentation in response to diverse unfolding stresses (28), although a higher temperature of 40 to 42°C is required to elicit a similar response. Comparison of the insoluble, detergent resistant protein fraction of wild type cultures with that of the GroESL depletion strain either during depletion or 36°C treatment in the presence of xylose revealed mild aggregation, or solubility changes, in a small number of proteins (Figure 1F). Together, these results indicate that proteostasis and growth are generally maintained in the early stages of GroESL insufficiency. Therefore, the division defect is likely to result from the misfolding of one or more specific proteins linked to the cell cycle that depend on an interaction with GroESL for functionality.

### Chromosome replication and cell cycle transcription continue during GroESL insufficiency

*C. crescentus* filamentation can result from perturbations in DNA replication, chromosome segregation, the cell cycle transcriptional program, or inhibition of the cell division machinery. To identify which stage(s) of the cell cycle GroESL folding is required for, we assessed the consequences of GroESL depletion on each of these processes. Measuring DNA content by flow cytometry revealed that GroESL-depleting cultures accumulate additional chromosomes, even late in depletion (Figure 2A), demonstrating that DNA replication continues when GroESL folding is insufficient. Consistent with this, multiple well-spaced origins of replication were distributed throughout the cytoplasm (Figure 2B), and the chromosomes were spread throughout the entirety of the cell body (Figure 2C). These data argue against a problem with chromosome segregation, which generally features chromosome-free spaces or mislocalized origins of replication (29).

**Figure 2:**
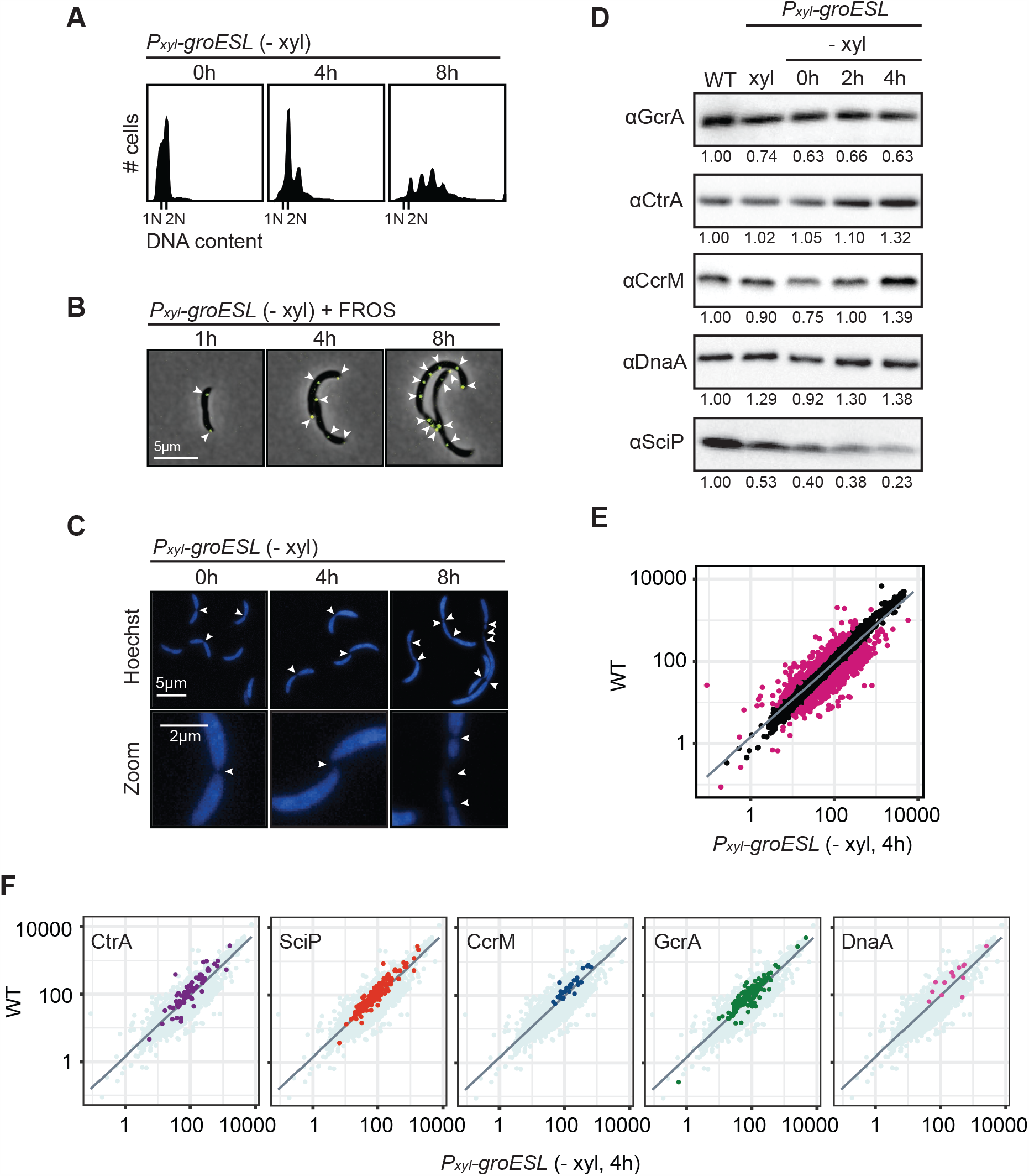
Chromosome replication and cell cycle transcription continue during GroESL insufficiency. (A) Flow cytometry profiles showing DNA content per cell at indicated time points of GroESL depletion. (B) Microscopy of fluorescently labeled origins of replication at indicated time points of GroESL depletion. A fluorescent reporter operator system (FROS) reporter construct bearing *ori::(tetO)n tetR-yfp* was used to mark origins of replication. Arrows indicate location of origin of replication foci. Images show YFP-phase merge. (C) Microscopy of cells at indicated time points during GroESL depletion, stained with Hoechst 33258 to visualize chromosomes. Arrows indicate gaps in staining associated with inter-chromosomal spaces. (D) Western blots of cell cycle regulators GcrA, CtrA, CcrM, DnaA, and SciP in wild type (WT) *C. crescentus* and GroESL depletion strain (*P*_*xyl*_*-groESL*), grown either in non-depleting conditions (+ xyl) or for the indicated time periods in depleting conditions (−xyl). Quantification of band intensities represent an average of three biological replicates. (E) RNAseq analysis showing normalized expression values for *C. crescentus* transcriptome of wild type versus GroESL depletion at four hours. Genes with a fold change less than −2 or greater than 2 are represented by deep pink points. Line represents smoothed conditional mean of data. (F) Plots as in (E) with the genes belonging to the regulons of GcrA (34), CtrA (36), CcrM (32), DnaA (35), and SciP (33) highlighted.

The transcriptional circuit driving the cell cycle is poised to halt at the appearance of many stress inputs (28, 30, 31), therefore we assessed GroESL-depleted cells for the presence of the major cell cycle regulators CtrA, DnaA, CcrM, GcrA and SciP, which drive the cell cycle-dependent transcriptional program in *C. crescentus* (32–36). CtrA, DnaA, GcrA and CcrM remained at near wild type levels during the four hours of GroESL depletion when the filamentation phenotype emerges, while SciP levels showed a reduction during this time frame (Figure 2D). It is possible that this reduction of SciP is caused by its increased degradation by the protease Lon, which is known to degrade SciP. Like other components of the proteostasis network (12, 15), we found Lon levels to increase in GroESL-depleted cells, offering a potential explanation for this observation (Supplemental Figure 1).

To more directly test if loss of GroESL-mediated folding affects cell cycle-regulated transcription in *Caulobacter*, we performed RNAseq analysis comparing wild type *C. crescentus* transcription with that during early GroESL depletion (Figure 2E). This analysis showed that the transcriptional regulons controlled by all five cell cycle regulators, including SciP, remained largely unchanged when GroESL folding is reduced (Figure 2F). Together these results show that DNA replication, chromosome segregation, and the cell cycle-dependent transcriptional program are not markedly affected by a reduction in available GroESL. Therefore, one or more proteins of the cell division apparatus may be specifically sensitive to the availability of GroESL, and mediate the filamentation phenotype of GroESL depletion.

### Loss of GroESL is associated with changes in solubility of division and PG synthesis proteins

To identify candidate division-linked proteins whose folding is perturbed by reduced GroESL, we utilized a quantitative proteomics approach using isobaric tandem mass tag (TMT) mass spectrometry to identify proteins enriched in the insoluble, detergent resistant fraction of cultures in early GroESL depletion (Figure 3A). We identified 630 proteins whose presence in the insoluble fraction was significantly different between wild type and GroESL depletion, including 167 proteins with abundances increased at least 1.5-fold (p < 0.05) (Supplemental Table 2). The best predictor of *E. coli* GroESL client proteins is a specific physicochemical signature (37), part of which is the presence of specific structural folds, therefore we analysed the folds present in our identified population of enriched insoluble proteins. As in *E. coli* (18, 19, 38), we found the TIM beta/alpha barrel fold (c.1) to be over-represented in proteins enriched in the *C. crescentus* GroESL-depleted insoluble fraction (Figure 3B). This enrichment of the c.1 fold is specific to GroESL depletion, as analysis of fold prevalence in the insoluble fraction of heat-stressed *C. crescentus* revealed other fold classes to be more prevalent in this condition (Supplemental Figure 2) (39). Among the proteins enriched in the insoluble fraction of GroESL-depleted cells, we identified five essential proteins that are linked to cell division and that function in the cell envelope; FzlA, FtsA, MurA, MurG, and DapA, as well as KidO, a non-essential oxidoreductase with a TIM beta/alpha barrel fold (Figure 3A) (40, 41). FzlA, FtsA and KidO are proteins that directly interact with FtsZ, the major structural component of the *Caulobacter* divisome (40–44), while MurA, MurG and DapA are part of the PG biosynthesis pathway (45–48), which is critical for maintaining the cell envelope during normal growth and building the new poles during division.

**Figure 3:**
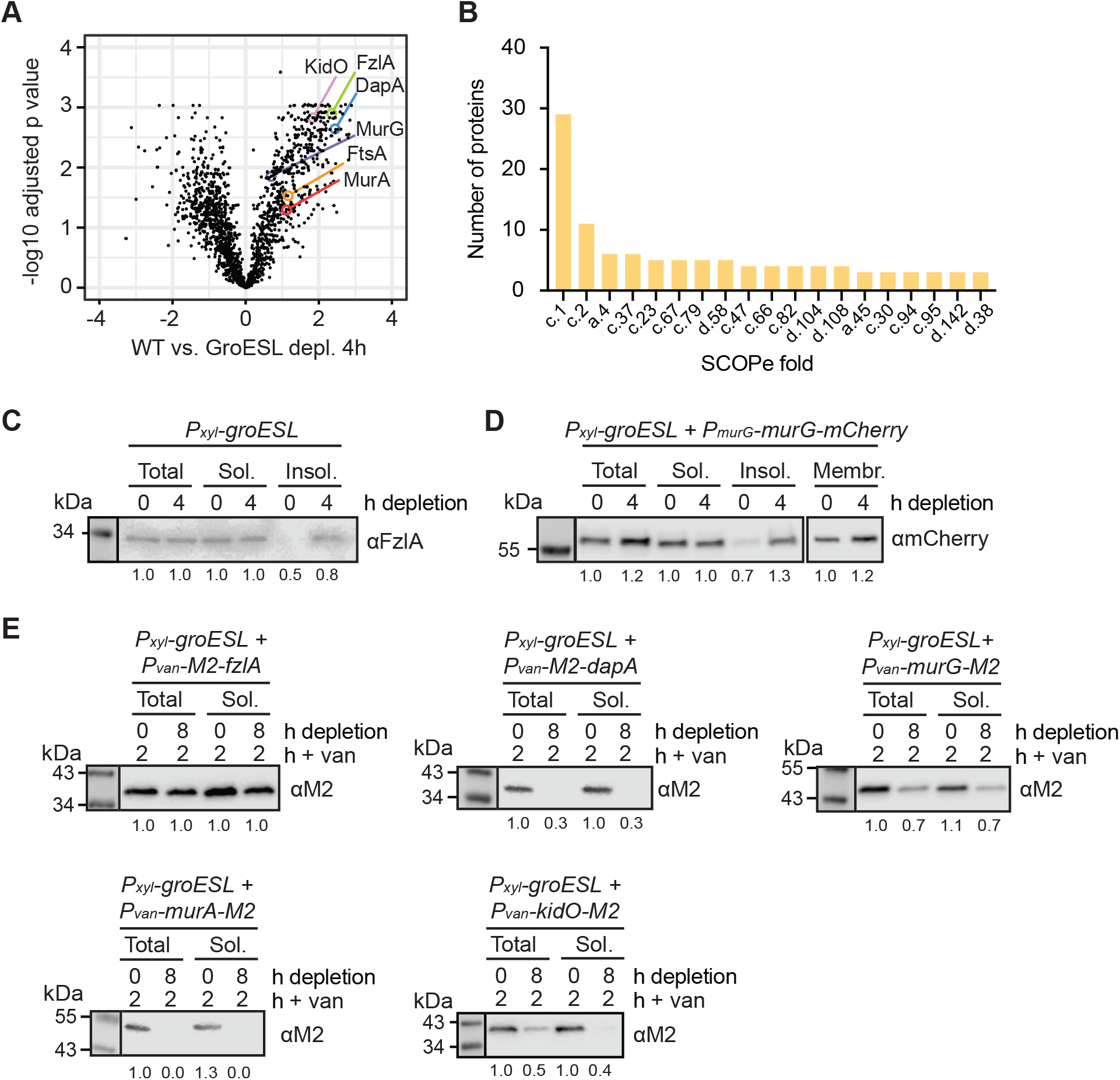
Loss of GroESL is associated with changes in division and cell wall synthesis protein solubility. (A) Changes in detergent-resistant insoluble fractions between wild type cultures and cultures depleted of GroESL for four hours, identified by quantitative proteomics. Volcano plot shows significance (log adjusted P value, calculated using linear model analysis) versus log fold change WT vs. GroESL depletion 4h. Identified PG synthesis and cell division proteins are indicated. (B) Enrichment of structural folds (SCOPe classification) in proteins of the detergent-resistant insoluble fraction of cultures depleted of GroESL for four hours. Number of proteins indicates the absolute number of proteins identified with the indicated fold ID. (C) Western blot of native FzlA abundance in cell lysate (total), soluble (sol.) and insoluble (insol.) cellular fractions of the GroESL depletion strain (*P*_*xyl*_*-groESL)*. Samples were taken from cultures where GroESL was not depleted (0h) as well as cultures where GroESL was depleted for four hours. Quantification of band intensities represents an average of three biological replicates. (D) Western blot of native MurG-mCherry abundance in cell lysate (total), soluble (sol.), insoluble (insol.), and membrane (membr.) cellular fractions of the GroESL depletion strain (*P*_*xyl*_*-groESL)*. Samples were taken from cultures where GroESL was not depleted (0h) as well as cultures where GroESL was depleted for four hours. Quantification of band intensities represents an average of 3 biological replicates. (E) Solubility of *de novo* synthesized M2-FzlA, M2-DapA, MurG-M2, MurA-M2 and KidO-M2 in cells depleted of GroESL (8h). GroESL was depleted for 0h or 6h prior to induction of M2-FzlA, M2-DapA, MurG-M2, MurA-M2, and KidO-M2 for 2h from the vanillate-inducible promoter (*P*_*van*_). Cultures were harvested and isolated into cell lysate (total) or soluble (sol.) fractions and immunoblotted. Quantification of band intensities represents an average of 3 biological replicates.

To validate the mass spectrometry results, we assessed the solubility of native FzlA and a MurG-mCherry fusion, integrated at the native chromosomal locus, when GroESL availability was reduced (Figure 3C, 3D). This confirmed that these proteins are significantly enriched in the insoluble fraction when GroESL was depleted (Figure 3C, 3D). Notably, a large proportion of both FzlA and MurG-mCherry was present in the soluble fractions. We reasoned that this could be either due to the majority of the protein achieving a folded and soluble state before GroESL levels become limiting, or because most of the newly produced protein can correctly fold in the absence of GroESL. To discriminate between these possibilities, we assessed more directly how GroESL availability affects *de novo* production of the candidate proteins. For this, we tagged FzlA, KidO, DapA, MurA, and MurG with a small M2 tag, induced the expression of these fusion proteins for two hours in either GroESL-depleted (6 hours) or non-depleted conditions, and then quantified their abundance in the total and soluble protein fractions (Figure 3E). We attempted to tag FtsA to include in this analysis, however as FtsA does not tolerate tags at either terminus, and tagging this protein has been demonstrated to alter its stability and function (49), we could not include it in our analysis. Our data show that similar amounts of M2-FzlA were present in the soluble and total protein fractions of GroESL-depleted and non-depleted cultures (Figure 3E), indicating that FzlA can be produced and accumulate in a soluble state with reduced levels of GroESL. By contrast, the M2-DapA, MurG-M2, MurA-M2, and KidO-M2 fusions were only present at high levels when GroESL levels were sufficient for viability (Figure 3E). In particular, *de novo* synthesized DapA and MurA were not tolerated in cells lacking GroESL (Figure 3E), therefore the accumulation of these proteins is strictly dependent on GroESL availability. We were however able to detect an enrichment in insoluble, native DapA and MurA in early GroESL depletion (Figure 3A), indicating that synthesis of these proteins during insufficient GroESL folding results in production of insoluble protein, which is then degraded. Similar behaviour is observed for several obligate *E. coli* GroESL substrates, including DapA (19, 20, 47). Our data suggest that DapA, MurA, MurG and KidO could be GroESL clients in *C. crescentus*, as their accumulation depends on the presence of GroESL. As soluble FzlA was produced and accumulated in GroESL-depleted cultures, we conclude that this protein is unlikely to be an obligate GroESL client, but do not yet exclude a contribution to the cell division defect of GroESL-depleted cells.

### GroESL folding supports PG biosynthesis through MurG, MurA, and DapA

DapA, MurA, and MurG are all part of the PG biosynthetic pathway, which functions to build and maintain the outer structure of growing cells, including the new cell poles during division (16). We first focussed on this group of proteins and investigated the relationship between PG biosynthesis and GroESL folding. First, we sought additional support for our data that the divisome-associated protein MurG may require an interaction with GroESL to reach a functional state, and determined the localization of MurG-mCherry during GroESL depletion. We found that MurG-mCherry formed multiple foci along the length of the cell (Figure 4A), which may be indicative of aggregation (39), or alternatively, as MurG function is associated with FtsZ and the Z-ring (26, 45, 48), its association with partially assembled or partially functional divisome components. Importantly, we observed that in a subpopulation of cells (30%) MurG-mCherry formed polar foci (Figure 4A, 4B). These polar MurG-mCherry foci did not occur in non-depleting conditions, suggesting that they are caused by reduced GroESL availability. As condensation of *Caulobacter* FtsZ, and therefore the divisome, is inhibited at the poles (50), and furthermore as the *Caulobacter* poles are stable regions where new PG is not inserted (45), this observation is consistent with MurG-mCherry clustering in a non-functional state.

**Figure 4:**
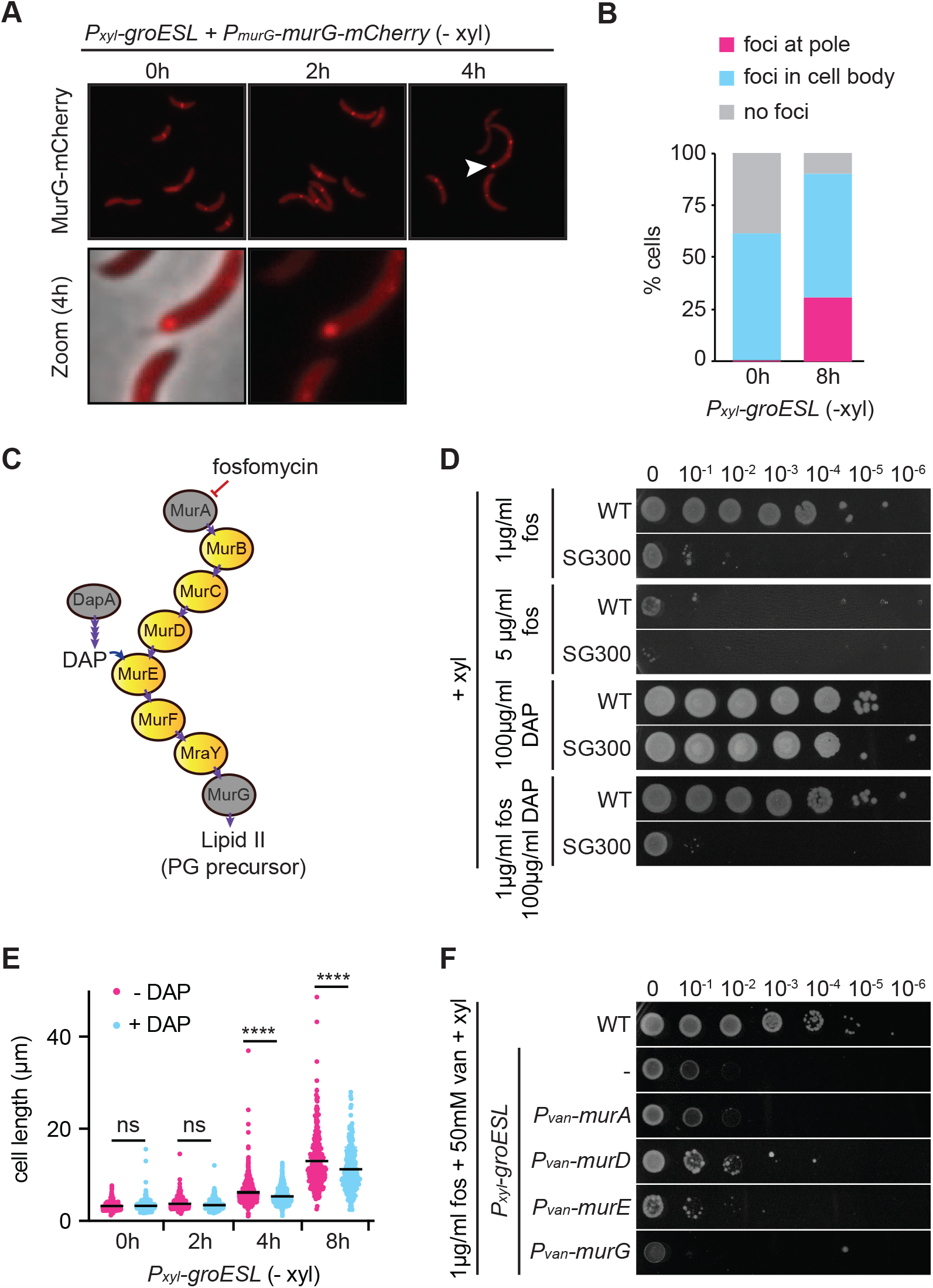
GroESL folding supports PG biosynthesis through MurG, MurA, and DapA. (A) Microscopy of MurG-mCherry localization during early GroESL depletion (0 to 4h). Representative images are shown, white arrow marks polar MurG-mCherry localization, magnified in lower panels. (B) Quantification of population location of MurG-mCherry localization patterns before (0h) and after eight hours of GroESL depletion (n < 339, graph is average of biological duplicates). Foci at pole; cell contains at least one focus that is located in the extreme polar region, foci in cell body; cell contains foci but not located at the pole, no foci; cell contains only diffuse signal. (C) Diagram of PG biosynthetic pathway in *C. crescentus*. Proteins identified in Figure 3A are highlighted in grey. Important metabolites (DAP, Lipid II) are indicated where they appear in the pathway, as well as where fosfomycin acts to inhibit MurA. Purple arrows indicate enzymatic reactions. (D) Spot assay of wild type and GroESL depletion strain (*P*_*xyl*_*-groESL)* in the presence of fosfomycin (fos) and/or DAP. Xylose was included in all agar plates. Images are representative of 3 biological replicates. (E) Quantification of population cell lengths (n < 244 each population) determined by phase contrast microscopy of the GroESL depletion strain (*P*_*xyl*_*-groESL)* during depletion (-xyl) in the presence or absence of 100µg/ml DAP (****, p < 0.0001). (F) Spot assay of wild type and derivatives of the GroESL depletion strain (*P*_*xyl*_*-groESL)* harboring chromosomally-encoded, inducible genes encoding the PG biosynthetic pathway proteins MurA-M2, MurD-M2, MurE-M2 or MurG-M2 when treated with fosfomycin (fos). Xylose was included in all agar plates to support GroESL expression, and vanillate was included to induce the expression of PG biosynthesis proteins. The GroESL depletion strain (*P*_*xyl*_*-groESL*) without integrated plasmids is included as a control (-). Images are representative of 3 biological replicates.

PG precursors are built in the cytoplasm through the sequential action of a series of enzymes, including MurA and MurG, and metabolites for the pathway are supplied by DapA (Figure 4C). The first committed step of PG biosynthesis requires the activity of MurA, which is targeted by the antibiotic fosfomycin (51, 52). To assess the stability of PG biosynthesis when GroESL folding is reduced, we determined the sensitivity of the GroESL depletion strain, grown in non-depleting conditions, to fosfomycin. Interestingly, the GroESL depletion strain demonstrated hypersensitivity towards this antibiotic (Figure 4D), indicating that PG biosynthesis is highly sensitive to changes in GroESL availability. In *E. coli*, DapA is an obligate GroESL substrate that catalyses the formation of 4-hydroxy-tetrahydrodipicolinate, a precursor of meso-diaminopimelate (DAP) that is required for normal PG synthesis (47). Addition of DAP to the growth medium prevents lysis due to DapA degradation in GroESL-depleted *E. coli* (47), therefore we supplemented fosfomycin-treated *C. crescentus* cultures with DAP in an attempt to bypass the essential function of this protein (Figure 4D). DAP supplementation was unable to restore fosfomycin resistance (Figure 4D), therefore we also tested the effect of DAP on GroESL-depleting cultures in the absence of the antibiotic. *C. crescentus* supplemented with DAP still became filamentous during GroESL depletion, however the cells were slightly shorter in length, indicating some improvement in the phenotype (Figure 4E, Supplemental Figure 3). These data are consistent with the finding that DapA is not the only protein of the PG biosynthetic pathway that has solubility changes when GroESL levels become limiting, but that the proteins MurA and MurG may also require GroESL-mediated folding.

To address the possible interaction of other PG biosynthetic pathway enzymes with GroESL, we attempted to increase the activity of these proteins by increasing their gene copy number and therefore expression levels, a method that has been used to investigate the contribution of specific clients to the phenotype of GroESL depletion in *E. coli* (14). Increased expression of these proteins did not improve fosfomycin sensitivity (Figure 4F), suggesting again that the PG biosynthesis pathway has multiple points of interaction with GroESL. We additionally tested the involvement of MreB, which organizes PG insertion in *Caulobacter* and is classified as a Class II, non-obligate substrate of *E. coli* GroESL (19, 48), and also as a client of DnaK (53). However, increased expression of the actin homologue MreB did not rescue PG hypersensitivity, and the GroESL depletion strain was not more sensitive to the MreB inhibitor A22 than wild type at the concentrations tested (Supplemental Figure 4). Together, our results indicate that GroESL supports the folding and solubility of several proteins of the PG biosynthesis pathway, including MurG, MurA and DapA. Decreased GroESL folding capacity results in reduced functionality of this pathway, and consequently increased fosfomycin sensitivity.

### The Z-ring stalls shortly after GroESL levels begin to decline

In addition to proteins required for PG biosynthesis, we identified FzlA, FtsA, and KidO as being enriched in the insoluble fraction of GroESL-depleted cells (Figure 3A). All three of these proteins interact with FtsZ, and FzlA and FtsA provide essential regulation of FtsZ polymer formation, and consequently its function in coordinating cell division (42, 44, 54, 55). Therefore, we determined the effects of GroESL depletion on the formation and function of the Z-ring. We first assessed the condensation of FtsZ during GroESL depletion using a merodiploid FtsZ-eYFP fusion reporter (Figure 5A) (50). We found that before significant GroESL-mediated cell length changes occur during GroESL depletion, the Z-ring was present at midcell in a larger proportion of the population than in actively dividing cells (Figure 5B). By 4h depletion, multiple FtsZ foci were present in disorganized locations along the cell length, with no obvious bias in positioning other than that FtsZ foci remained excluded from the poles (Figure 5A, 5B). Therefore, when GroESL levels are reduced, FtsZ polymerizes and condenses but stalls before division is complete, with many of the Z-rings assembled within two hours of GroESL depletion failing to achieve division. The ability to divide is lost asynchronously, as division is observed to occur in some cells later in depletion, suggesting that the remaining chaperonin may occasionally provide enough folding of the required division protein(s) (Supplemental Movie 1). Collectively, these results indicate that a stalling of the Z-ring immediately precedes the cell length changes observed during early GroESL depletion, suggesting that misfolding of an FtsZ-interacting protein is the primary driver of the cell division defect.

**Figure 5:**
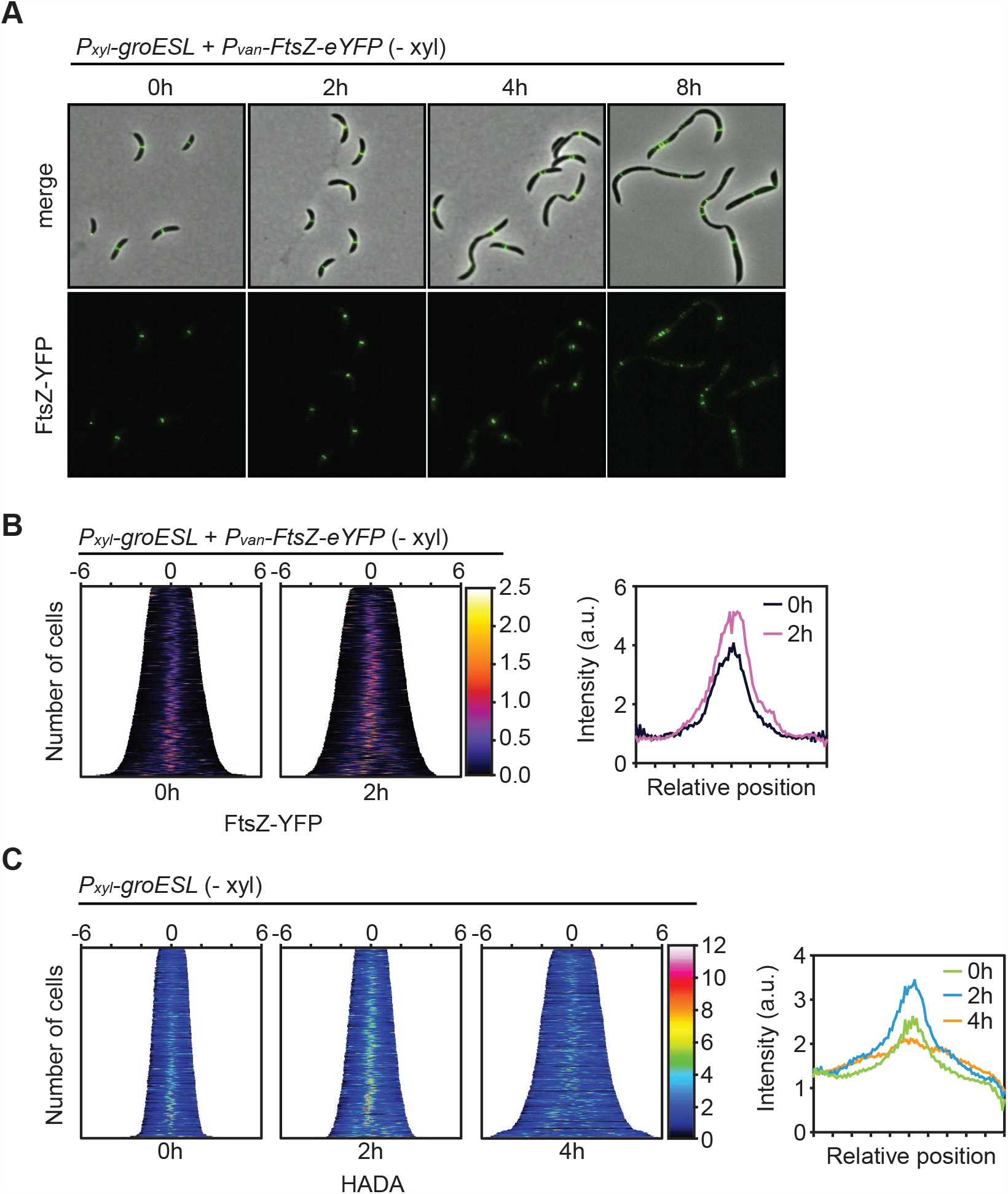
The Z-ring stalls shortly after GroESL levels decline. (A) Microscopy of FtsZ-eYFP localization in the GroESL depletion strain during depletion (-xyl). FtsZ-eYFP was expressed from the vanillate-inducible promoter (*P*_*van*_*-ftsZ-eYFP*) for two hours prior to imaging each time point of GroESL depletion. Representative micrographs are shown. (B) Demographs showing fluorescent signal profiles of FtsZ-eYFP in early GroESL depletion (0-2h), organized by cell length (n < 308 each population). Fluorescent profiles are organized by cell length. (C) Demographs of population fluorescence intensity profiles of HADA stain (n <334 each population). Cultures were depleted of GroESL for the indicated time periods and exposed to a short pulse (2 min) of HADA prior to fixation and imaging. Population intensity profiles are organized by cell length.

To confirm that FtsZ stalling at midcell is not an artifact of the fluorescent fusion construct, we further assessed Z-ring formation using a fluorescent-D-amino acid (HADA) (Figure 5C), which marks the active PG incorporation at midcell that is coordinated by FtsZ, as well as that coordinated by the elongasome (56). In agreement with the FtsZ fluorescent fusion, we observed that almost all of the population exhibited a bright midcell focus of PG incorporation at 2h GroESL depletion, in contrast to actively dividing cells (Figure 5C). Additionally, foci became disorganized and were found along the cell length at later time points (Figure 5C), indicating that the Z-ring continues to coordinate PG insertion while being unable to complete cell division. Furthermore, we did not observe FtsZ foci or foci of PG insertion to occur at the poles with HADA staining, consistent with our hypothesis that some of the MurG present in GroESL-depleted cells is clustering in a non-functional, insoluble state.

### GroESL folding regulates FtsZ ring function not through FzlA, but FtsA

Because both FzlA and FtsA are critical for regulating FtsZ dynamics (42, 44, 55), we hypothesized that incorrect or insufficient folding of either, or both, of these proteins may lead to the observed changes in FtsZ behavior at the early stages of GroESL depletion. To evaluate the effects of FzlA on Z-ring function, we made use of a previously established *ΔfzlA* suppressor strain (54), in which a point mutation in FtsW (A246T) compensates for loss of the essential function of FzlA. However, depletion of GroESL in the *ΔfzlA* suppressor strain did not improve or delay the filamentation phenotype (Figure 6A, B), and the development of the characteristic irregularly spaced constrictions was still observed (Figure 6A). This result suggests that FzlA does not contribute significantly to the cell division defect of GroESL depletion. While not essential for *Caulobacter* division (57), we also tested the involvement of the ABC transporter and FtsZ-interacting protein FtsE (Figure 6A, 6B), due to its involvement in the GroESL depletion phenotype of *E. coli* (19, 21). However, as with FzlA, the GroESL phenotype developed similarly in a *C. crescentus* strain lacking *ftsE* (Figure 6A, 6B), confirming FtsE does not mediate the phenotype in this organism.

**Figure 6:**
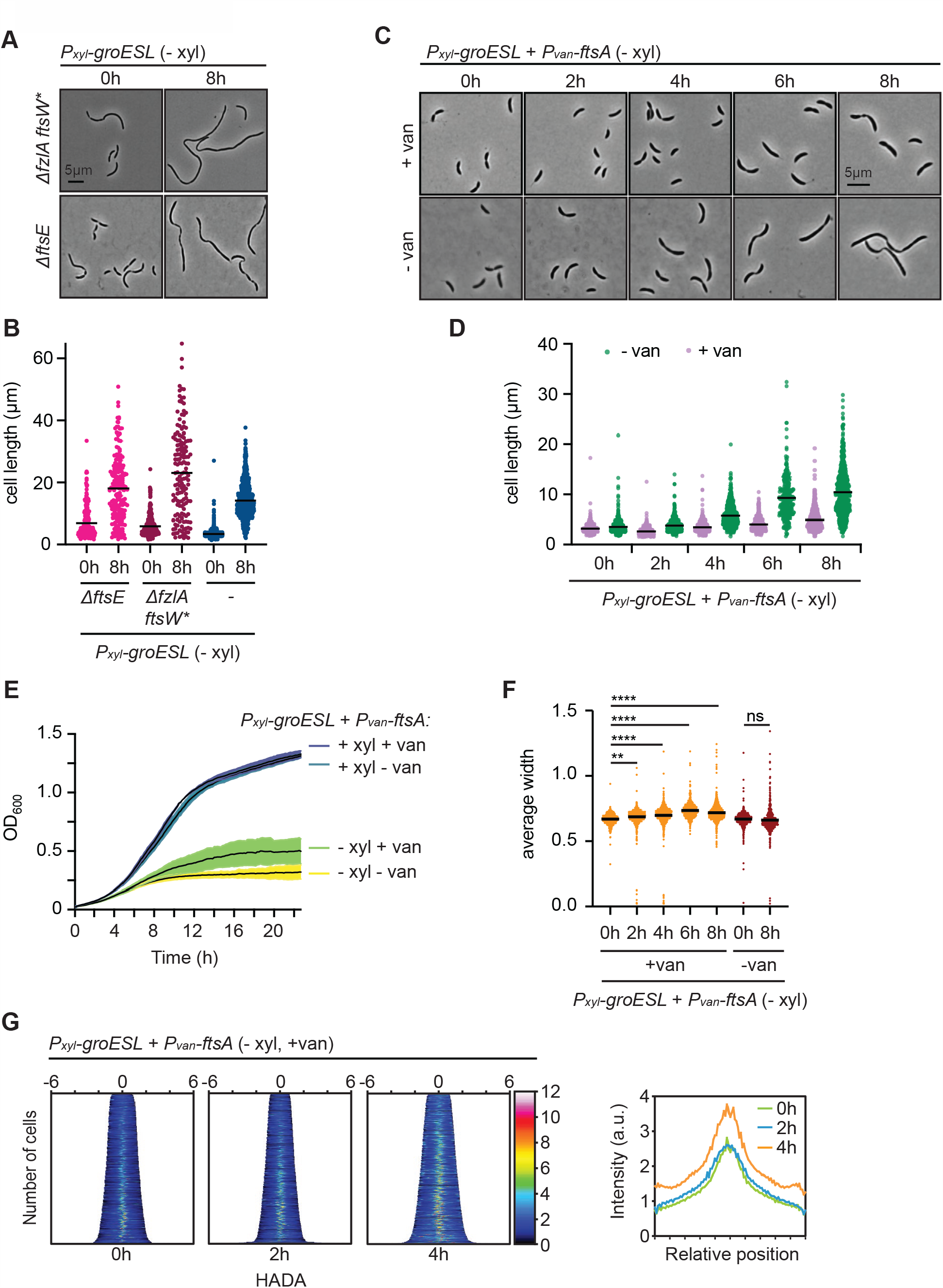
GroESL folding regulates FtsZ ring function through FtsA, not FzlA or FtsE. (A) Microscopy of strains lacking FzlA (*ΔfzlAftsW\**) or FtsE (*ΔftsE*) before and eight hours after GroESL depletion. (B) Quantification of population cell lengths of the strains shown in (A), compared with population cell lengths of the parent GroESL depletion strain (shown in blue). (C) Microscopy of GroESL depletion with induced expression of *ftsA* from a second chromosomal locus. Vanillate-dependent *ftsA* expression was induced at the onset of GroESL depletion (0h). Microscopy images of isogenic cultures during GroESL depletion, grown without the addition of vanillate (-van) are shown for comparison. (D) Quantification of population cell lengths of (C) (n < 299, each population). Isogenic cultures were grown with (+van) or without (-van) the addition of vanillate and population cell lengths quantified. (E) Growth curve assessing biosynthetic capacity of the GroESL depletion strain (*P*_*xyl*_*-groESL)* producing additional FtsA. Isogenic cultures were grown in the presence or absence of xylose (+/-xyl) and the presence of absence of vanillate (+/- van) to determine growth effects. (F) Quantification of population mean widths of cells producing extra FtsA. Cell widths were measured in populations from (C) and (D). ANOVA was used to determine population differences (**, p < 0.0013, ****, p < 0.0001). (G) Population fluorescence intensity profiles of HADA stain for populations of GroESL-depleted cultures producing extra FtsA (n < 502, each population). Cultures were depleted of GroESL for the indicated time periods with vanillate-dependent expression of FtsA induced at the onset of depletion (0h). At the indicated time points cultures were exposed to a short pulse (2 min) of HADA prior to fixation and imaging. Population intensity profiles are organized by cell length.

We next investigated whether impaired functioning of FtsA contributes to the filamentation observed in GroESL depletion. In particular, FtsA seemed a promising candidate, as FtsA depletion results in filamentous cells with shallow, irregularly spaced constrictions that resemble the phenotype of GroESL depletion (58). As FtsA cannot be suppressed or deleted, we sought to evaluate if increased FtsA production could reduce or delay the effects of insufficient GroESL folding and alleviate the GroESL depletion phenotype (Figure 6C). Strikingly, when FtsA was produced from an additional chromosomal locus we observed a significant delay in the development of filamentation during GroESL depletion (Figure 6C, 6D). Cells were shorter and contained fewer constrictions than in GroESL-depleted cultures without additional FtsA (Figure 6C, 6D), suggesting additional division events had occurred. These observations are consistent with increased FtsA production being able to compensate for a reduction in the ability to efficiently fold FtsA. It is important to note that the FtsA expression levels in this genetic context did not lead to the filamentation phenotypes observed with strong overexpression of FtsA in wild type *C. crescentus* (Supplemental Figure 5) (59). Growth analysis revealed that production of extra FtsA improved growth capacity (Figure 6E), thus excluding the possibility that the shorter cells observed during GroESL depletion in the presence of additional FtsA were due to growth arrest or a decrease in growth rate. Finally, the shorter cells were also wider than those observed during GroESL depletion in the absence of additional FtsA (Figure 6F). As cell thickening is associated with defects in PG biosynthesis, we hypothesize that in the event of restoring sufficient FtsA, PG instability is the dominant phenotype that emerges due to GroESL depletion.

To evaluate the impact of providing extra FtsA on the function of FtsZ during early GroESL depletion (Figure 5C), we again used HADA staining. A delay in Z-ring stalling was observed (Figure 6G), where the proportion of newly divided cells without a midcell focus of PG incorporation was maintained for an additional 2 hours, or at least one additional population doubling (Figure 6G vs. Figure 5C), during which growth rate was maintained. Collectively these experiments illustrate that GroESL is necessary to support normal Z-ring function during division, and while GroESL insufficiency results in solubility changes for FzlA and FtsA, it is FtsA function that is most sensitive to the folding capacity of the chaperonin. Furthermore, our data demonstrate that the interaction between FtsE and GroESL, required for division in *E. coli*, is not conserved in *C. crescentus*, which instead tunes cell division to chaperone availability through an actin protein-chaperonin interaction.

## Discussion

Chaperonins are highly conserved folding machines that provide essential protein folding across all kingdoms of life. Critical functions of chaperonins range from helping bacteria to build peptidoglycan (47) to supporting chloroplast and mitochondrial function in eukaryotes (60, 61), and information from prokaryotic systems has helped to inform exploration of human chaperonins (62). In this present work we expand on how chaperonin function is integrated into bacterial physiology by exploring GroESL function in the alphaproteobacterium *Caulobacter crescentus*. We find that the integration of GroESL into the processes of cell division and synthesis of the cell envelope is conserved amongst different groups of bacteria, however this integration occurs via distinct points of interaction (Figure 7). In *C. crescentus*, GroESL folding is required to support PG biosynthesis via MurG, MurA, and DapA, but is most critically required to support cell division through an interaction with FtsA. By linking a chaperonin to these processes, stress-responsive protein folding capacity is intimately connected to both cell envelope synthesis and cell division in *Caulobacter*.

**Figure 7:**
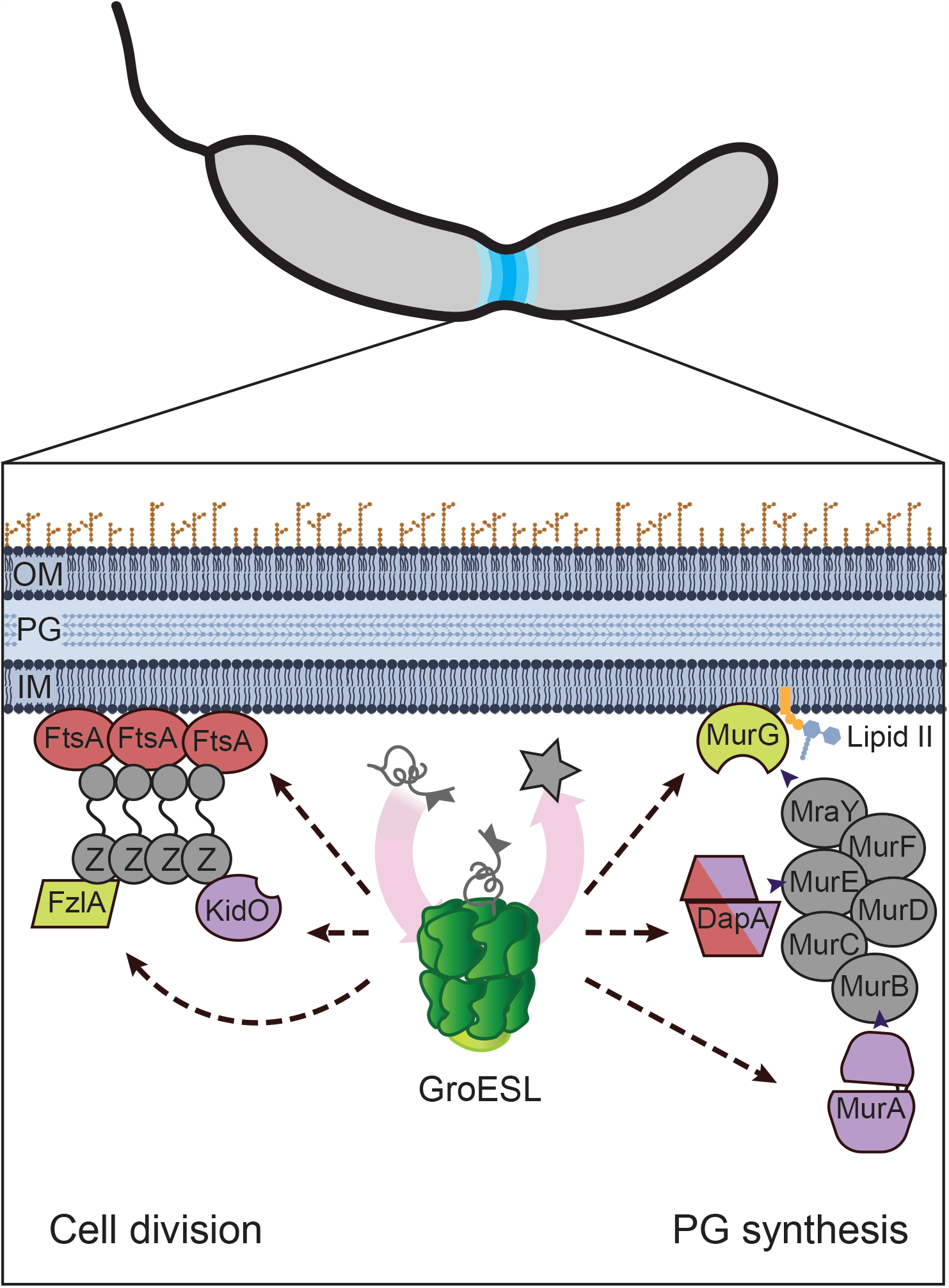
Model of GroESL folding supporting *Caulobacter crescentus* cell envelope biosynthesis and cell division. GroESL supports the function of several division proteins and peptidoglycan biosynthetic enzymes during *C. crescentus* division, and is most critically required for supporting FtsA. The chaperonin GroESL is a foldase that assists in moving client proteins from folding intermediates to native folded proteins (star). GroESL folding supports solubility of FtsA, FzlA, and KidO in the divisome, and MurG, DapA, and MurA in the PG biosynthetic pathway (metabolite flow from candidate GroESL client proteins indicated by arrowheads). Proteins in red (FtsA, DapA) are able to temporarily rescue the GroESL depletion filamentation phenotype if provided in excess or if their function is bypassed. FtsA interacts with FtsZ (Z) to anchor FtsZ filaments to the membrane and regulate its dynamics during division. Proteins in purple (DapA, MurA, KidO) are degraded if synthesized in the absence of sufficient GroESL folding. Proteins in green exhibit altered localization. Proteins in grey do not decrease solubility during GroESL depletion. Dashed lines represent interactions that may proceed through an as of yet unidentified intermediate. OM; outer membrane, PG; peptidoglycan, IM; inner membrane. Membrane and PG images created with Biorender (Biorender.com).

Our study has shown that chaperonin folding is indispensable for PG synthesis in *C. crescentus*, and has identified several new interactions between PG biosynthetic proteins and GroESL. This is the first description of MurA being linked to GroESL folding, though an interaction with DnaKJE has previously been established (53). Our data suggests that MurG solubility and localization may also respond to GroESL-mediated folding (Figure 4A). MurG has been shown to act as a scaffold for PG biosynthesis in *Bordetella pertussis* and *Thermotoga maritima* (63, 64) and could provide a similar function in *C. crescentus*, though it remains unclear if the changes we observe for MurG are due to absence of a direct interaction with GroESL, or perhaps loss of an upstream signal required for PG biosynthetic subcomplex assembly. Depletion of PG precursors, which may occur through a reduction in pathway protein function, results in filamentation to conserve limited resources and prevent over-investment in the intensive process of building new cell poles (51). During periods of proteotoxic stress and high refolding demand, titration of the chaperonin away from synthetic processes could provide a way to postpone cell division and focus on survival. Interaction of both DnaKJE and GroESL with unfolded proteins is known to regulate the heat shock response to this end (5), and DnaKJE availability during stress is integrated into the cell cycle as an indirect regulator of DNA replication initiation (31). It remains to be discovered how PG synthetic protein folding and abundance are prioritized during stress. Newly discovered accessory factors, such as the holdase CnoX (65), may hold the key to how client proteins are presented to GroESL, and which processes are protected during high unfolding demand. Discerning these interactions will be important to understanding how organisms balance growth and division when surviving stress.

Our study has identified the bacterial actin homologue FtsA as a protein that is particularly sensitive to GroESL availability. FtsA was among the proteins that showed increased insolubility in the absence of GroESL (Figure 3A), and although its enrichment in the insoluble fraction was mild, producing extra FtsA alone was able to delay the development of filamentation as chaperonin levels declined (Figure 6D), to a greater extent than other interventions (Figure 4E). Work in *E. coli* has shown that increasing the expression of other GroESL client proteins (or those that feed into the client protein function) can also temporarily compensate for reduced levels of GroESL during depletion (14, 21), by providing a reserve pool of folded client protein to draw on. Our data linking FtsA function with GroESL is particularly striking, as a major role of the eukaryotic chaperonin TRiC is to perform folding of eukaryotic actin (66, 67), yet bacterial FtsA has not been identified previously as an interactor of GroESL or DnaKJE (19, 53). Therefore, our findings raise questions on the conservation of the relationship between actin proteins and chaperonins. As FtsA is also a highly conserved and crucial protein in diverse bacteria, it will be important to determine the relationship between actin homologues and GroESL in other organisms, including clarifying this relationship in *E. coli*. Our work has excluded a role for the *E. coli* obligate GroESL client protein FtsE in *C. crescentus* (19, 21), suggesting that different organisms have evolved separate links between GroESL and cell division. As GroESL is thought to support evolutionary plasticity in metabolic enzymes (68), it is an open question if the chaperonin might permit similar flexibility in cell division proteins, a question with consequences for resistance to current and future antimicrobials that target the cell envelope and cell division.

## Materials and Methods

### Strains and plasmids

The strains and plasmids used in this study are listed in Supplemental Table 3.

### Bacterial growth conditions

All *C. crescentus* strains were routinely cultured at 30°C, unless otherwise indicated, in liquid PYE media while shaking at 200 rpm. If necessary, the following media supplements were added to the following final concentrations: 0.3% xylose, 0.2% glucose, 25µg/ml spectinomycin, 5µg/ml kanamycin, 0.625µg/ml gentamycin, 500mM vanillate. Cultures were regularly diluted to keep them in mid-log phase. In GroESL depletion experiments, cells were washed three times with PYE free of media supplements by centrifugation (6000 *xg*, 4 min) before resuspension in medium lacking xylose inducer. Growth on solid PYE media was performed in the presence of the following supplement concentrations: 0.3% xylose, 0.2% glucose, 500mM vanillate, 5µg/ml gentamycin, 25µg/ml kanamycin, 400µg/ml spectinomycin. Transductions were performed using ϕCr30 as described previously (69). *E. coli* was grown for cloning purposes in LB supplemented with antibiotics as necessary at 37°C.

### Spot assays

Spot assays were performed with cultures maintained in log phase for three hours and diluted to an OD_600_ of ∼0.2. Tenfold serial dilutions of this culture were prepared and 2µL of each dilution was spotted and dried onto a fresh agar plate.

### Growth curves

For growth curve experiments, cultures maintained in log phase for 3 hours were diluted to an OD_600_ of ∼0.05 and 200µL diluted culture was added to 96-well plates. Measurement was performed every ten minutes at 30°C with culture aeration in a Tecan Spark for 24h. Three biological replicates were performed for all growth curve measurements, with three technical replicates for each sample.

### Western blotting

Cell pellets were harvested by centrifugation and resuspended in Laemmli buffer normalized to OD_600_ measurement, followed by heating at 70°C for 10min. Protein extracts were loaded on 4-20% stain-free SDS-PAGE gels and subjected to electrophoresis before activation and transfer to a nitrocellulose membrane. Successful transfer and equal loading were verified by 2,2,2-trichloroethanol visualization prior to blotting. Specific proteins were detected using the following primary antibody dilutions: anti-CtrA; 1:5,000 (kindly provided by MT Laub), anti-GcrA; 1:4,000 (70), anti-CcrM; 1:5,000 (71), anti-DnaA; 1:5,000 (72), anti-SciP; 1:2,000 (33), anti-Lon; 1:10,000 (kind gift from RT Sauer), anti-GroEL; 1:10,000 (8), and the commercially available anti-M2 1:1,000 (Sigma). HRP-conjugated secondary antibody raised against rabbit or mouse was used at a 1:5,000 dilution, and SuperSignal Femto West reagent was used for signal detection using a Licor Odyssey. Images were processed and quantified using Fiji.

### Microscopy and image analysis

For cell length analysis, samples were fixed in 1% formaldehyde and spotted on 1% agarose pads. A final concentration of 2µg/ml Hoechst 33258 was used to stain fixed cells by incubating 25 minutes in the dark prior to mounting. HADA staining was performed on ethanol-fixed cells as in (73). For live cell imaging, including all instances of fluorescent protein imaging, the microscope housing was heated to 30°C and live cells were spotted on 1% agar PYE pads containing xylose, glucose, or vanillate as necessary.

Imaging was performed on a Nikon Ti-Eclipse microscope equipped with a 100X objective and Zyla 4.2 Plus camera, and at least ten independent frames of each sample were collected using Nikon Image Elements AR software. Image stacks were imported into Fiji and background of fluorescent images was subtracted prior to segmentation using MicrobeJ (74). In all images, segmentation was manually checked prior to exporting data. Unless otherwise indicated, ANOVA analysis (including adjustment for multiple comparisons where necessary) was performed to derive statistical significance of morphological changes using GraphPad Prism 8 software.

### Flow cytometry

Samples of *C. crescentus* cultures grown as indicated were fixed in a final concentration of 70% ethanol. Cells were pelleted and washed in 50mM sodium citrate buffer containing 2µg/ml RNase, and incubated overnight at 50°C. A final concentration of 2.5µg/ml SYTOX green was used to stain 1:10 dilutions of the RNA-digested samples immediately prior to processing by a BD Biosciences LSR-Fortessa flow cytometer. Data were analysed and histograms prepared with FlowJo.

### Subcellular fractionation

Isolation of the detergent-resistant insoluble fraction was adapted from (39), as follows. Log phase cultures were harvested at the indicated time points or conditions and pelleted at 7000xg for 10min at 4°C. Cells were washed once in buffer I (50mM Tris-HCl pH 8.0, 150mM NaCl) and frozen at −80°C. Pellets were resuspended in buffer I supplemented with 12 U/ml benzonase and disrupted by sonication (10 cycles of 30s on, 30s off at 50% amplitude in a QSonica sonicator). Cellular debris was removed from the lysate by centrifugation at 5000xg for 10min at 4°C and removing supernatant, and repeating this step. Protein concentration of lysate was determined by Nanodrop. To separate soluble and insoluble fractions, lysate was centrifuged at 20,000xg for 20min at 4°C. The insoluble fraction was washed in buffer I, resuspended by one cycle of sonication, and pelleted again, followed by incubation with 1% Triton X-100 for 1h with regular vortexing. The insoluble fraction was pelleted again and washed an additional two times before resuspension in Laemmli buffer. Dilution in Laemmli buffer was normalized according to lysate protein concentration, with insoluble fractions were concentrated 20X to account for the lower relative abundance of this fraction. Membrane fractions were prepared separately as in (75).

### RNA sequencing

RNA of bacterial cultures was extracted using the RNeasy mini kit (Qiagen), and RNA sequencing performed by GENEWIZ (South Plainfield, NJ). Gene expression data are available at the Gene Expression Omnibus repository: GSE162320.

### Mass spectrometry

The insoluble, detergent resistant fraction of cultures was harvested and prepared in biological duplicates according to the protocol described above. Protein digestion, TMT10plex isobaric labelling and mass spectrometry were performed at the Clinical Proteomics Mass Spectrometry facility (Karolinska Institute, Karolinska University Hospital, Science for Life Laboratory). To determine differential abundance in the insoluble fractions, linear model analysis was performed as in (76). Only significantly changed protein abundances (p > 0.05) were considered for further analysis as described in the text. For analysis of SCOP folds, fold identity was predicted from amino acid sequence using the SUPERFAMILY 2 database (77, 78).

## Acknowledgements

We thank Dr. Suely Gomes for providing the SG300 strain and GroEL antibody, Dr. Erin Goley for the kind gifts of strains, plasmids, FzlA antibody, and helpful discussions, and Dr. Patrick Viollier for providing CcrM and GcrA antibodies. We also thank the Clinical Proteomics Mass Spectrometry facility and National Bioinformatics Infrastructure Sweden (NBIS, SciLifeLab) for assistance with collecting and analysing the proteomics data as well as members of the Jonas group for suggestions and comments. The study was financially supported by the Swedish Foundation for Strategic Research (FFL15-0005), the Swedish Research Council (2016-03300), and funding from the Strategic Research Area (SFO) program distributed through Stockholm University.

## Supplemental Information

**Supplemental Figure 1:**
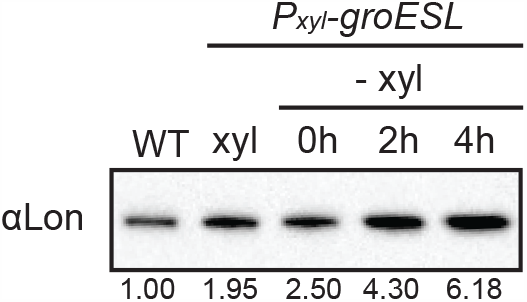
The Lon protease is upregulated upon GroESL depletion. Western blot showing Lon abundance in wild type (WT) *C. crescentus* and the GroESL depletion strain (*P*_*xyl*_*-groESL*), grown either in non-depleting conditions (+ xyl) or for 0, 2 and 4 hours in depleting conditions (-xyl). Quantification of band intensities represent an average of three biological replicates.

**Supplemental Figure 2:**
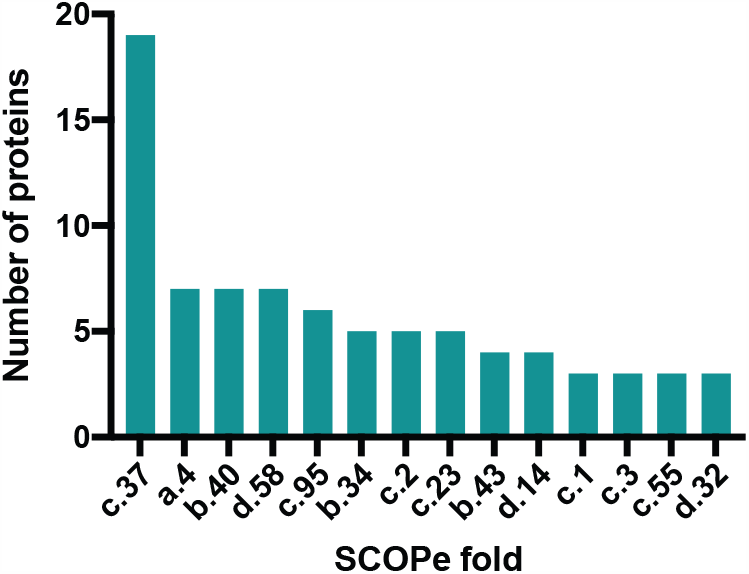
Fold distribution of aggregated proteins in heat stress. Enrichment of structural folds (SCOPe classification) in proteins of the detergent-resistant insoluble fraction of wild type *C. crescentus* cultures exposed to 45°C for one hour. Number of proteins indicates the absolute number of proteins identified with the indicated fold ID. Data set was used from (39).

**Supplemental Figure 3:**
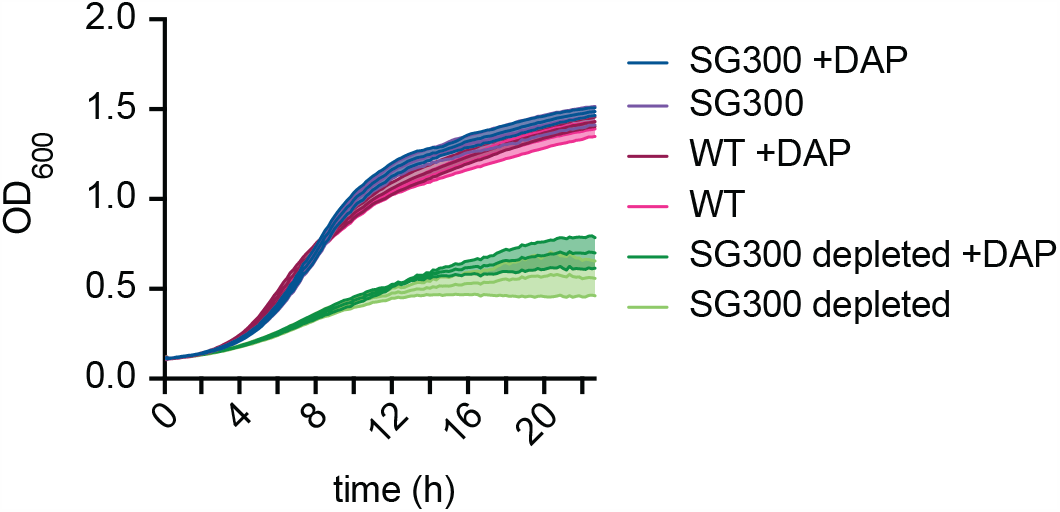
Growth curve during DAP supplementation in GroESL depletion. Growth curve assessing biosynthetic capacity of wild type and GroESL depletion strains in the presence and absence of 100µg/ml DAP. Cultures were prepared at an OD of 0.1 and depleted prior to adding to the plate containing the appropriate additives where necessary.

**Supplemental Figure 4:**
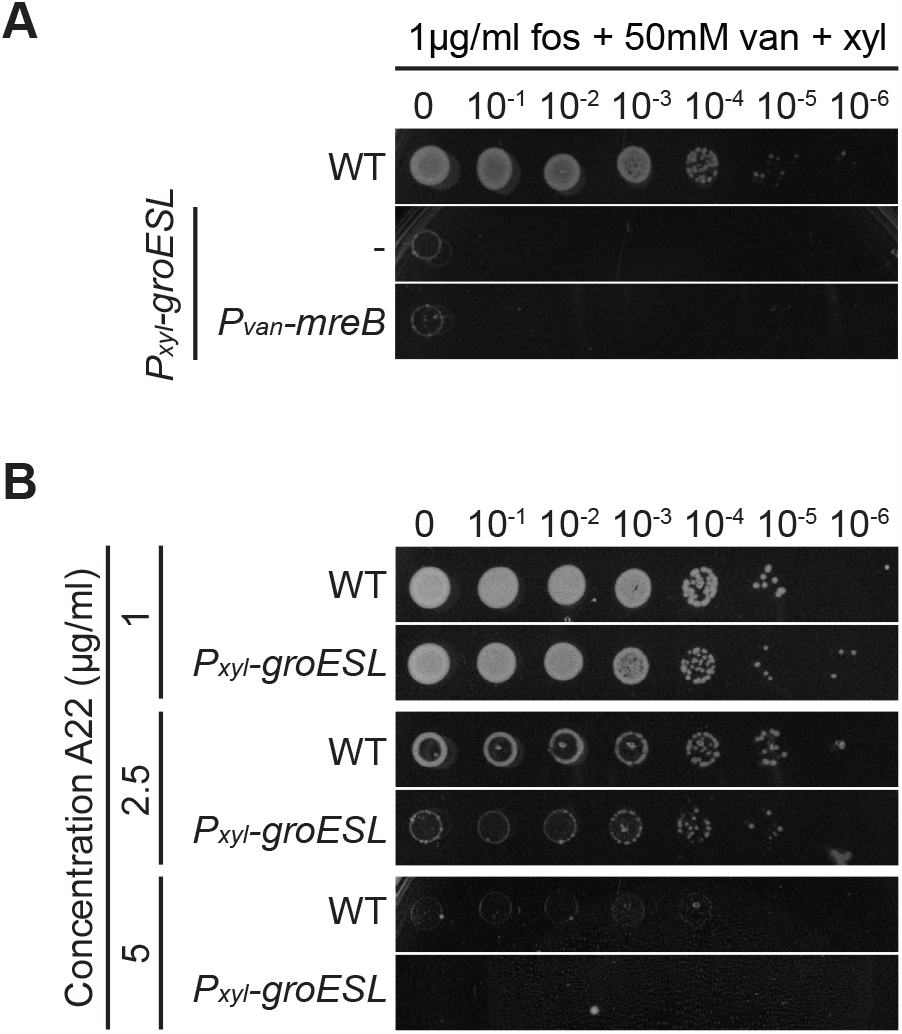
MreB does not mediate the GroESL-dependent PG defect. (A) Spot assay of wild type and derivative of the GroESL depletion strain (*P*_*xyl*_*-groESL)* harboring a chromosomally-encoded, inducible M2-MreB when treated with fosfomycin (fos). Xylose was included in all agar plates to support GroESL expression, and vanillate was included to induce the expression of MreB. The GroESL depletion strain (*P*_*xyl*_*-groESL*) without integrated plasmids is included as a control (-). Images are representative of 3 biological replicates. (B) Spot assay of wild type and GroESL depletion strain (*P*_*xyl*_*-groESL)* in the presence of the MreB inhibitor A22. Xylose was included in all agar plates. Images are representative of 3 biological replicates.

**Supplemental Figure 5:**
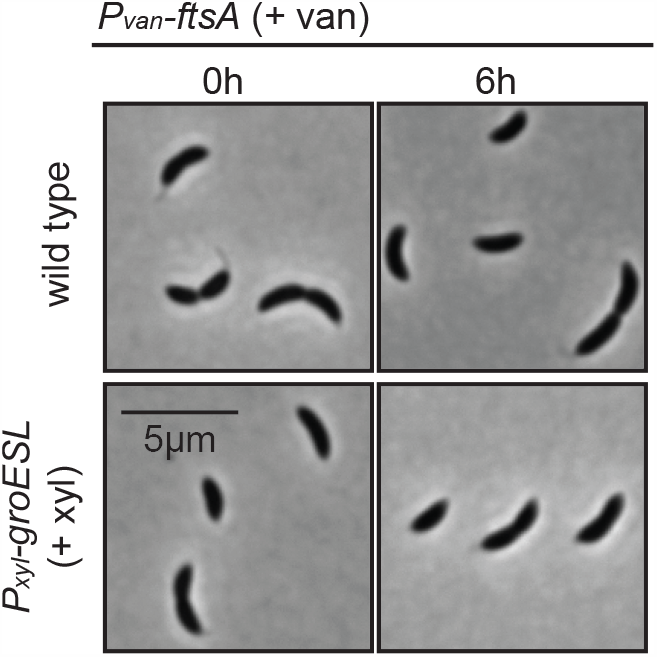
Phenotype of FtsA expression from a second chromosomal locus. Microscopy of induced expression of *ftsA* from a second chromosomal locus in wild type and GroESL depletion strain (+xyl). Microscopy images are shown from exponentially growing cultures prior to *ftsA* induction (0h), and after vanillate-dependent *ftsA* expression was induced and maintained in exponentially growing cultures for 6h.

**Supplemental Table 1: RNAseq data**

**Supplemental Table 2: Proteomics data**

**Supplemental Table 3: Strain and plasmid list**

**Supplemental Movie 1: Dynamics of FtsZ condensation during GroESL depletion**.

## References

1. Balchin D, Hayer-Hartl M, Hartl FU. 2020. Recent advances in understanding catalysis of protein folding by molecular chaperones. FEBS Lett 594:2770–2781.

2. Anfinsen CB, Scheraga HA. 1975. Experimental and theoretical aspects of protein folding. Adv Protein Chem 29:205–300.

3. Dahl J-U, Gray MJ, Jakob U. 2015. Protein quality control under oxidative stress conditions. J Mol Biol 427:1549–1563.

4. Tamás MJ, Fauvet B, Christen P, Goloubinoff P. 2018. Misfolding and aggregation of nascent proteins: a novel mode of toxic cadmium action in vivo. Curr Genet 64:177–181.

5. Roncarati D, Scarlato V. 2017. Regulation of heat-shock genes in bacteria: from signal sensing to gene expression output. FEMS Microbiol Rev 41:549–574.

6. Schramm FD, Schroeder K, Jonas K. 2020. Protein aggregation in bacteria. FEMS Microbiol Rev 44:54–72.

7. Hayer-Hartl M, Bracher A, Hartl FU. 2016. The GroEL-GroES Chaperonin Machine: A Nano-Cage for Protein Folding. Trends Biochem Sci 41:62–76.

8. Baldini RL, Avedissian M, Gomes SL. 1998. The CIRCE element and its putative repressor control cell cycle expression of the Caulobacter crescentus groESL operon. J Bacteriol 180:1632–1641.

9. Lund PA. 2009. Multiple chaperonins in bacteria – why so many? FEMS Microbiol Rev 33:785–800.

10. Glass JI, Lefkowitz EJ, Glass JS, Heiner CR, Chen EY, Cassell GH. 2000. The complete sequence of the mucosal pathogen Ureaplasma urealyticum. Nature 407:757–762.

11. Chowdhury N, Kingston JJ, Whitaker WB, Carpenter MR, Cohen A, Boyd EF. 2014. Sequence and expression divergence of an ancient duplication of the chaperonin groESEL operon in Vibrio species. Microbiology 160:1953–1963.

12. ACA Da Silva, Simão RCG, Susin MF, Baldini RL, Avedissian M, Gomes SL. 2003. Downregulation of the heat shock response is independent of DnaK and σ32 levels in Caulobacter crescentus: Heat shock response regulation in Caulobacter. Mol Microbiol 49:541–553.

13. Lemos JA, Luzardo Y, Burne RA. 2007. Physiologic effects of forced down-regulation of dnaK and groEL expression in Streptococcus mutans. J Bacteriol 189:1582–1588.

14. Masters M, Blakely G, Coulson A, McLennan N, Yerko V, Acord J. 2009. Protein folding in Escherichia coli: the chaperonin GroE and its substrates. Res Microbiol 160:267–277.

15. Susin MF, Baldini RL, Gueiros-Filho F, Gomes SL. 2006. GroES/GroEL and DnaK/DnaJ have distinct roles in stress responses and during cell cycle progression in Caulobacter crescentus. J Bacteriol 188:8044–8053.

16. Egan AJF, Errington J, Vollmer W. 2020. Regulation of peptidoglycan synthesis and remodelling. Nat Rev Microbiol https://doi.org/10.1038/s41579-020-0366-3.

17. Haeusser DP, Margolin W. 2016. Splitsville: structural and functional insights into the dynamic bacterial Z ring. Nat Rev Microbiol 14:305–319.

18. Fujiwara K, Ishihama Y, Nakahigashi K, Soga T, Taguchi H. 2010. A systematic survey of in vivo obligate chaperonin-dependent substrates. EMBO J 29:1552–1564.

19. Kerner MJ, Naylor DJ, Ishihama Y, Maier T, Chang H-C, Stines AP, Georgopoulos C, Frishman D, Hayer-Hartl M, Mann M, Hartl FU. 2005. Proteome-wide analysis of chaperonin-dependent protein folding in Escherichia coli. Cell 122:209–220.

20. Niwa T, Fujiwara K, Taguchi H. 2016. Identification of novel in vivo obligate GroEL/ES substrates based on data from a cell-free proteomics approach. FEBS Lett 590:251–257.

21. Fujiwara K, Taguchi H. 2007. Filamentous morphology in GroE-depleted Escherichia coli induced by impaired folding of FtsE. J Bacteriol 189:5860–5866.

22. Govezensky D, Greener T, Segal G, Zamir A. 1991. Involvement of GroEL in nif gene regulation and nitrogenase assembly. J Bacteriol 173:6339–6346.

23. Ogawa J, Long SR. 1995. The Rhizobium meliloti groELc locus is required for regulation of early nod genes by the transcription activator NodD. Genes Dev 9:714–729.

24. Ojha A, Anand M, Bhatt A, Kremer L, Jacobs WR, Hatfull GF. 2005. GroEL1: a dedicated chaperone involved in mycolic acid biosynthesis during biofilm formation in mycobacteria. Cell 123:861–873.

25. Curtis PD, Brun YV. 2010. Getting in the loop: regulation of development in Caulobacter crescentus. Microbiol Mol Biol Rev MMBR 74:13–41.

26. Goley ED, Yeh Y-C, Hong S-H, Fero MJ, Abeliuk E, McAdams HH, Shapiro L. 2011. Assembly of the Caulobacter cell division machine. Mol Microbiol 80:1680–1698.

27. Zielińska A, Billini M, Möll A, Kremer K, Briegel A, Izquierdo Martinez A, Jensen GJ, Thanbichler M. 2017. LytM factors affect the recruitment of autolysins to the cell division site in Caulobacter crescentus: The autolytic machinery of C. crescentus. Mol Microbiol 106:419–438.

28. Heinrich K, Sobetzko P, Jonas K. 2016. A Kinase-Phosphatase Switch Transduces Environmental Information into a Bacterial Cell Cycle Circuit. PLoS Genet 12:e1006522.

29. Ward D, Newton A. 1997. Requirement of topoisomerase IV parC and parE genes for cell cycle progression and developmental regulation in Caulobacter crescentus. Mol Microbiol 26:897–910.

30. Jonas K. 2014. To divide or not to divide: control of the bacterial cell cycle by environmental cues. Curr Opin Microbiol 18:54–60.

31. Jonas K, Liu J, Chien P, Laub MT. 2013. Proteotoxic stress induces a cell-cycle arrest by stimulating Lon to degrade the replication initiator DnaA. Cell 154:623–636.

32. Gonzalez D, Kozdon JB, McAdams HH, Shapiro L, Collier J. 2014. The functions of DNA methylation by CcrM in Caulobacter crescentus: a global approach. Nucleic Acids Res 42:3720–3735.

33. Gora KG, Tsokos CG, Chen YE, Srinivasan BS, Perchuk BS, Laub MT. 2010. A cell-type-specific protein-protein interaction modulates transcriptional activity of a master regulator in Caulobacter crescentus. Mol Cell 39:455–467.

34. Haakonsen DL, Yuan AH, Laub MT. 2015. The bacterial cell cycle regulator GcrA is a σ70 cofactor that drives gene expression from a subset of methylated promoters. Genes Dev 29:2272–2286.

35. Hottes AK, Shapiro L, McAdams HH. 2005. DnaA coordinates replication initiation and cell cycle transcription in Caulobacter crescentus. Mol Microbiol 58:1340–1353.

36. Laub MT, Chen SL, Shapiro L, McAdams HH. 2002. Genes directly controlled by CtrA, a master regulator of the Caulobacter cell cycle. Proc Natl Acad Sci U S A 99:4632–4637.

37. Azia A, Unger R, Horovitz A. 2012. What distinguishes GroEL substrates from other Escherichia coli proteins?: GroEL substrates. FEBS J 279:543–550.

38. Niwa T, Ying B-W, Saito K, Jin W, Takada S, Ueda T, Taguchi H. 2009. Bimodal protein solubility distribution revealed by an aggregation analysis of the entire ensemble of Escherichia coli proteins. Proc Natl Acad Sci U S A 106:4201–4206.

39. Schramm FD, Schroeder K, Alvelid J, Testa I, Jonas K. 2019. Growth-driven displacement of protein aggregates along the cell length ensures partitioning to both daughter cells in Caulobacter crescentus. Mol Microbiol 111:1430–1448.

40. Beaufay F, Coppine J, Mayard A, Laloux G, De Bolle X, Hallez R. 2015. A NAD-dependent glutamate dehydrogenase coordinates metabolism with cell division in Caulobacter crescentus. EMBO J 34:1786–1800.

41. Radhakrishnan SK, Pritchard S, Viollier PH. 2010. Coupling prokaryotic cell fate and division control with a bifunctional and oscillating oxidoreductase homolog. Dev Cell 18:90–101.

42. Barrows JM, Sundararajan K, Bhargava A, Goley ED. 2020. FtsA Regulates Z-Ring Morphology and Cell Wall Metabolism in an FtsZ C-Terminal Linker-Dependent Manner in Caulobacter crescentus. J Bacteriol 202.

43. Din N, Quardokus EM, Sackett MJ, Brun YV. 1998. Dominant C-terminal deletions of FtsZ that affect its ability to localize in Caulobacter and its interaction with FtsA. Mol Microbiol 27:1051–1063.

44. Goley ED, Dye NA, Werner JN, Gitai Z, Shapiro L. 2010. Imaging-Based Identification of a Critical Regulator of FtsZ Protofilament Curvature in Caulobacter. Mol Cell 39:975–987.

45. Aaron M, Charbon G, Lam H, Schwarz H, Vollmer W, Jacobs-Wagner C. 2007. The tubulin homologue FtsZ contributes to cell elongation by guiding cell wall precursor synthesis in Caulobacter crescentus. Mol Microbiol 64:938–952.

46. Barreteau H, Kovač A, Boniface A, Sova M, Gobec S, Blanot D. 2008. Cytoplasmic steps of peptidoglycan biosynthesis. FEMS Microbiol Rev 32:168–207.

47. McLennan N, Masters M. 1998. GroE is vital for cell-wall synthesis. Nature 392:139.

48. White CL, Kitich A, Gober JW. 2010. Positioning cell wall synthetic complexes by the bacterial morphogenetic proteins MreB and MreD: Control of cell shape in bacteria. Mol Microbiol 76:616–633.

49. Williams B, Bhat N, Chien P, Shapiro L. 2014. ClpXP and ClpAP proteolytic activity on divisome substrates is differentially regulated following the Caulobacter asymmetric cell division. Mol Microbiol 93:853–866.

50. Thanbichler M, Shapiro L. 2006. MipZ, a spatial regulator coordinating chromosome segregation with cell division in Caulobacter. Cell 126:147–162.

51. Harris LK, Theriot JA. 2016. Relative Rates of Surface and Volume Synthesis Set Bacterial Cell Size. Cell 165:1479–1492.

52. Kahan FM, Kahan JS, Cassidy PJ, Kropp H. 1974. The mechanism of action of fosfomycin (phosphonomycin). Ann N Y Acad Sci 235:364–386.

53. Calloni G, Chen T, Schermann SM, Chang H-C, Genevaux P, Agostini F, Tartaglia GG, Hayer-Hartl M, Hartl FU. 2012. DnaK functions as a central hub in the E. coli chaperone network. Cell Rep 1:251–264.

54. Lariviere PJ, Mahone CR, Santiago-Collazo G, Howell M, Daitch AK, Zeinert R, Chien P, Brown PJB, Goley ED. 2019. An Essential Regulator of Bacterial Division Links FtsZ to Cell Wall Synthase Activation. Curr Biol 29:1460-1470.e4.

55. Pichoff S, Lutkenhaus J. 2005. Tethering the Z ring to the membrane through a conserved membrane targeting sequence in FtsA: Membrane tethering of Z ring by FtsA. Mol Microbiol 55:1722–1734.

56. Kuru E, Hughes HV, Brown PJ, Hall E, Tekkam S, Cava F, de Pedro MA, Brun YV, VanNieuwenhze MS. 2012. In Situ probing of newly synthesized peptidoglycan in live bacteria with fluorescent D-amino acids. Angew Chem Int Ed Engl 51:12519–12523.

57. Meier EL, Daitch AK, Yao Q, Bhargava A, Jensen GJ, Goley ED. 2017. FtsEX-mediated regulation of the final stages of cell division reveals morphogenetic plasticity in Caulobacter crescentus. PLOS Genet 13:e1006999.

58. Martin ME, Trimble MJ, Brun YV. 2004. Cell cycle-dependent abundance, stability and localization of FtsA and FtsQ in Caulobacter crescentus: FtsA and FtsQ cell cycle regulation in C. crescentus. Mol Microbiol 54:60–74.

59. Sackett MJ, Kelly AJ, Brun YV. 1998. Ordered expression of ftsQA and ftsZ during the Caulobacter crescentus cell cycle. Mol Microbiol 28:421–434.

60. Christensen JH, Nielsen MN, Hansen J, Füchtbauer A, Füchtbauer E-M, West M, Corydon TJ, Gregersen N, Bross P. 2010. Inactivation of the hereditary spastic paraplegia-associated Hspd1 gene encoding the Hsp60 chaperone results in early embryonic lethality in mice. Cell Stress Chaperones 15:851–863.

61. Zhao Q, Liu C. 2017. Chloroplast Chaperonin: An Intricate Protein Folding Machine for Photosynthesis. Front Mol Biosci 4:98.

62. Conway de Macario E, Robb FT, Macario AJL. 2016. Prokaryotic Chaperonins as Experimental Models for Elucidating Structure-Function Abnormalities of Human Pathogenic Mutant Counterparts. Front Mol Biosci 3:84.

63. Favini-Stabile S, Contreras-Martel C, Thielens N, Dessen A. 2013. MreB and MurG as scaffolds for the cytoplasmic steps of peptidoglycan biosynthesis: Mur ligases interact with MreB and MurG. Environ Microbiol 15:3218–3228.

64. Laddomada F, Miyachiro MM, Jessop M, Patin D, Job V, Mengin-Lecreulx D, Le Roy A, Ebel C, Breyton C, Gutsche I, Dessen A. 2019. The MurG glycosyltransferase provides an oligomeric scaffold for the cytoplasmic steps of peptidoglycan biosynthesis in the human pathogen Bordetella pertussis. Sci Rep 9.

65. Goemans CV, Beaufay F, Arts IS, Agrebi R, Vertommen D, Collet J-F. 2018. The Chaperone and Redox Properties of CnoX Chaperedoxins Are Tailored to the Proteostatic Needs of Bacterial Species. mBio 9.

66. Balchin D, Miličić G, Strauss M, Hayer-Hartl M, Hartl FU. 2018. Pathway of Actin Folding Directed by the Eukaryotic Chaperonin TRiC. Cell 174:1507-1521.e16.

67. Gao Y, Thomas JO, Chow RL, Lee G-H, Cowan NJ. 1992. A cytoplasmic chaperonin that catalyzes β-actin folding. Cell 69:1043–1050.

68. Takemoto K, Niwa T, Taguchi H. 2011. Difference in the distribution pattern of substrate enzymes in the metabolic network of Escherichia coli, according to chaperonin requirement. BMC Syst Biol 5:98.

69. Ely B. 1991. Genetics of Caulobacter crescentus. Methods Enzymol 204:372–384.

70. Holtzendorff J, Hung D, Brende P, Reisenauer A, Viollier PH, McAdams HH, Shapiro L. 2004. Oscillating global regulators control the genetic circuit driving a bacterial cell cycle. Science 304:983–987.

71. Stephens C, Reisenauer A, Wright R, Shapiro L. 1996. A cell cycle-regulated bacterial DNA methyltransferase is essential for viability. Proc Natl Acad Sci 93:1210–1214.

72. Jonas K, Chen YE, Laub MT. 2011. Modularity of the Bacterial Cell Cycle Enables Independent Spatial and Temporal Control of DNA Replication. Curr Biol 21:1092–1101.

73. Kuru E, Tekkam S, Hall E, Brun YV, Van Nieuwenhze MS. 2015. Synthesis of fluorescent D-amino acids and their use for probing peptidoglycan synthesis and bacterial growth in situ. Nat Protoc 10:33–52.

74. Ducret A, Quardokus EM, Brun YV. 2016. MicrobeJ, a tool for high throughput bacterial cell detection and quantitative analysis. Nat Microbiol 1:16077.

75. Anwari K. 2012. Isolate and Sub-fractionate Cell Membranes from Caulobacter crescentus. BIO-Protoc 2.

76. Zhu Y, Orre LM, Zhou Tran Y, Mermelekas G, Johansson HJ, Malyutina A, Anders S, Lehtiö J. 2020. DEqMS: A Method for Accurate Variance Estimation in Differential Protein Expression Analysis. Mol Cell Proteomics 19:1047–1057.

77. Gough J, Karplus K, Hughey R, Chothia C. 2001. Assignment of homology to genome sequences using a library of hidden Markov models that represent all proteins of known structure. J Mol Biol 313:903–919.

78. Pandurangan AP, Stahlhacke J, Oates ME, Smithers B, Gough J. 2019. The SUPERFAMILY 2.0 database: a significant proteome update and a new webserver. Nucleic Acids Res 47:D490–D494.

